# A FLIC motility assay reveals myosin-6 coordination limited by actin filament buckling

**DOI:** 10.1101/068163

**Authors:** Agata K. Krenc, Jagoda J. Rokicka, Ronald S. Rock

## Abstract

Teams of myosin motors carry out intracellular transport and contract the actin cytoskeleton. To fully understand the behavior of multi-myosin ensembles we need to know the properties of individual myosins and the mode of interaction between them. Current models of the interactions within the myosin complex treat the actin filament as a stiff rod, not contributing to the regulation of collective myosin dynamics. Here, we present data suggesting that force transduction through the actin filament is an important element of interaction within myosin-6 ensembles *in vitro*. Multiple myosin-6s coordinate their steps if they are separated by a short (and therefore high-force bearing) segment of actin. The measurements were performed using Fluorescence Interference Contrast Microscopy (FLIC) to measure small changes in the height of fluorescently labeled actin. Using FLIC, we assign the positions of myosins in a gliding filament assay geometry and measure their attachment time to actin. We also identify actin segments that are buckled or under tension. We show that myosin-6 holds actin about 10 nm above the surface. However, due to asynchronous myosin stepping, frequent buckles up to about 60 nm high appear. The buckle lifetime decreases as the distance between the myosin-6s is reduced, a sign of inter-motor coordination. Our data are consistent with coordinated stepping of closely spaced myosins, but uncoordinated motility with widely separated myosins where buckles can form. These features would be expected to operate on myosins in the cell, where motor spacing may vary considerably depending on the target organelle.

**SIGNIFICANCE STATEMENT:** Myosins are molecular motors that carry out intracellular transport. Interactions between the myosins are crucial for understanding their function. Using Fluorescence Interference Contrast (FLIC) microscopy we characterized the interaction between multiple myosin-6 motors immobilized to the surface of a slide and pulling the same actin filament. Our results point towards coordination of myosin steps as a mechanism governing the behavior of a multi-myosin complex. We also demonstrated the unique application of FLIC microscopy for highly parallel identification and measurement of single myosin motors in a gliding filament format. These features of FLIC enable a robust study of collective myosin dynamics.

## INTRODUCTION

Myosins are molecular motors that hydrolyze ATP to power their movement along actin tracks [Howards 2001]. Multiple copies of class V (myosin-5) and class VI (myosin-6) myosins are associated with molecular cargos, suggesting that teams of processive motors govern intracellular transport (1–3). Over the last fifteen years, development of single molecule techniques enabled the detailed studies of biophysical properties of a single motor (4). However, a group of processive myosins can have very different properties from the individual myosins. The connection between the properties of a single myosin and a myosin assembly can be complex. How myosins behave when they are part of multi-motor assembly is an important question in the motor dynamics and cellular biology fields.

*In vitro*, an ensemble of motors typically demonstrates enhanced run-length and slower movement relative to a single motor (5–7). The critical role of collective motor behavior for the regulation of intracellular trafficking has been depicted by Efremov et. al (8). They showed that myosin 5–driven transport, but not kinesin-1-driven transport, was sensitive to motor density. The addition of myosins increased the velocity of a cargo, although the velocity was still below that of a single myosin. These studies highlight the importance of effective load-sharing for intracellular transport (8).

Myosin-6 is the only known myosin that walks towards the pointed end of actin (9). It was originally discovered in *Drosophila melanogaster* and has been shown to be important in early *Drosophila* development (10, 11). Myosin-6 works in a wide range of cellular roles including active transport and regulation of endocytosis (1). It also serves a structural or anchoring role, for example in maintaining separated stereocilia (12) or the maintenance of the correct morphology of the Golgi (13). The unique load-sensitivity of myosin-6 suggests a possible mechanism for its anchoring role at sub-saturating ATP conditions or in the presence of ADP (14). However, the load-sensitivity alone does not explain how myosin-6 could act as a vesicle transporter and anchor in the same cell, unless for each of these functions it experiences dramatically different load.

Here, we propose another mechanism of regulation of myosin-6 function, relying on its ability to effectively share the load between myosins carrying cargo along the same actin track. For a small cargo (e.g. endocytic vesicle) myosins are spaced close to each other, however for a large cargo (e.g., the Golgi apparatus) the myosins can be located further apart. Since actin is a semiflexible polymer capable of bending in a solution over a physiologically relevant range of lengths (15), the segment of actin can transduce a different amount of force for short and long separations. This could differentially regulate the activity of myosins, and they could be mechanically coupled or uncoupled from each other depending on the length of actin between them.

To address the question of myosin-6 collective behaviors as a function of the separation between the motors, we performed a series of experiments using fluorescence interference contrast microscopy (FLIC) (16, 17). FLIC measures the height of a fluorescently labeled actin filament above the surface in a gliding filament assay geometry. This allows for the detection of small changes in a filament contour line. The myosin attachments that tether actin to the surface are clearly visible in the assay. Therefore, using FLIC we can simultaneously observe the states of individual motors and the net product of their cooperation in a gliding filament assay. The data show that asynchronous myosin stepping leads to extensive actin buckling at intermediate myosin densities. However, as the density increases, the myosins coordinate their steps leading to short-lived buckles and decreased overall buckling.

## THE FLIC GLIDING FILAMENT ASSAY DESIGN

In the FLIC assay, we immobilized myosin on a silicon wafer, and then monitored the transport of fluorescently labeled actin filaments in a gliding filament assay. We stabilized the actin filaments with a fluorescent phalloidin at saturation, achieving uniform filament labeling. The schematic representation of the experimental set-up is depicted in Figure 1A. The principle of FLIC is the interference of light emitted by fluorophores near a reflective surface. The interference causes the observed fluorescence intensity to change as a function of a distance between the fluorophore and the reflective surface. To demonstrate that in FLIC the fluorescence intensity of a uniformly labeled actin is a reporter of its height above the reflective surface, we prepared several silicon wafers that differ in thickness of a transparent silicon oxide layer (see Materials and Methods). Static actin filaments, tethered to the surface as in Figure 1A, vary in fluorescence intensity with the thickness of the silicon oxide layer. These filament intensity data were used to create a calibration curve (Figure 1B), used later to determine the height of myosin-induced deformations. The peak of fluorescence observed in the FLIC calibration experiment corresponds to the height of ~80 nm above the surface, while the smaller heights result in dimmer fluorescence. This range and variation is consistent with previous reports (18–20).

**Figure 1.**
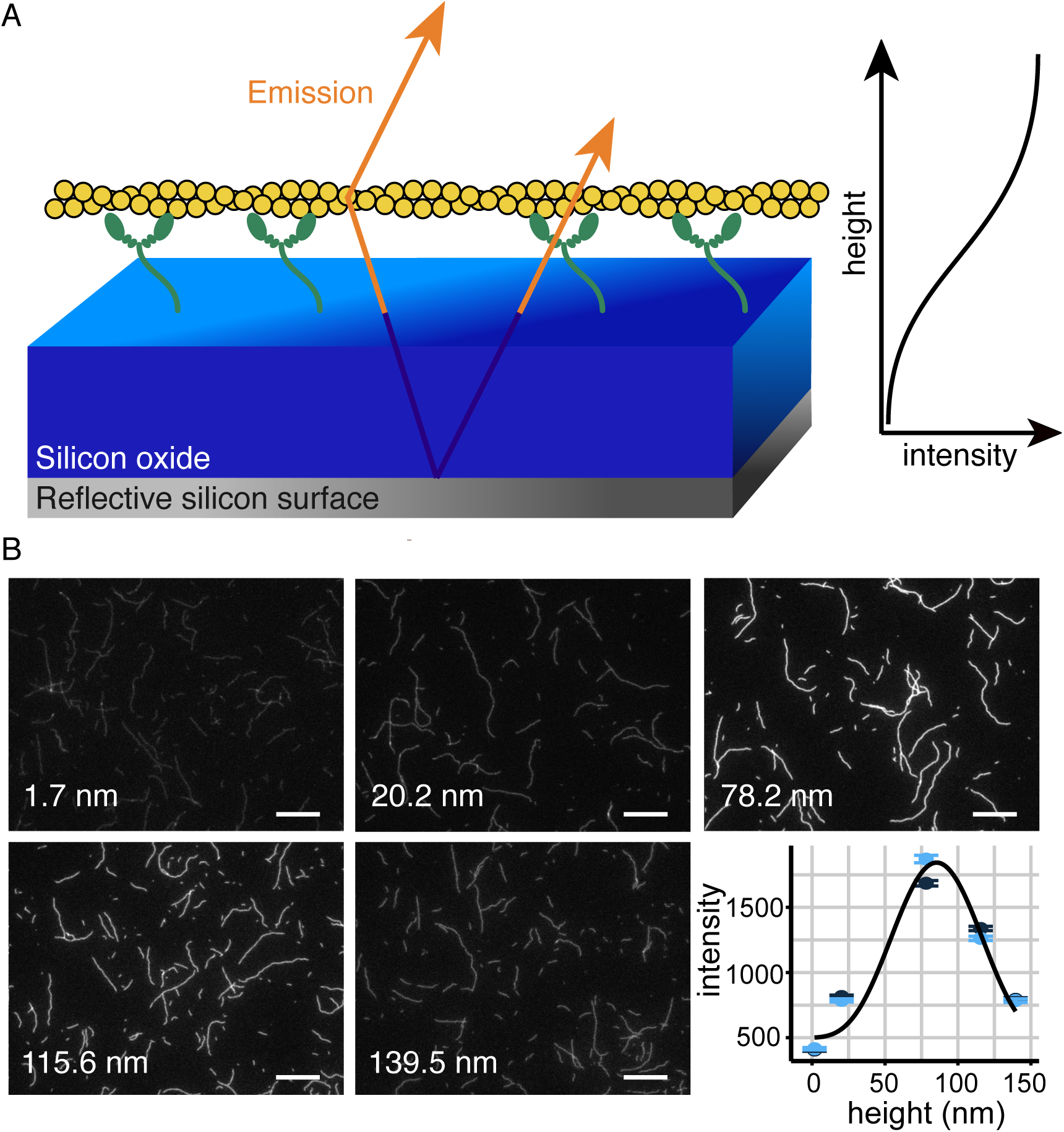
The FLIC assay measures distances between the fluorescently labeled actin and the Si/SiO_2_ surface. (A) The FLIC assay geometry. Myosins (green) are immobilized on the surface of a silicon wafer with a known thickness of a transparent silicon oxide layer (blue). Uniformly stained actin (yellow) adheres to the myosins. (B) FLIC calibration. Part of a single field of view from the microscope for each thickness of silicon oxide layer. We used the same preparation of actin for all oxide layer thicknesses shown. We applied myosin at 42 nM. Scale bar, 10 μm. The lower right panel shows the calibration curve, relating the fluorescence intensity to the height in nm. The data were fit to a model of FLIC intensity vs. height, black line (see SI). Black and blue dots represent two sets of measurements performed on the same day. Error bars show standard error of the mean in each measurement.

In the following sections of this manuscript the thickness of the silicon oxide layer is constant within anexperiment and the reported heights refer to the distance between the fluorescently labeled filament and the surface to which the myosins are immobilized (Figure 2A).

**Figure 2.**
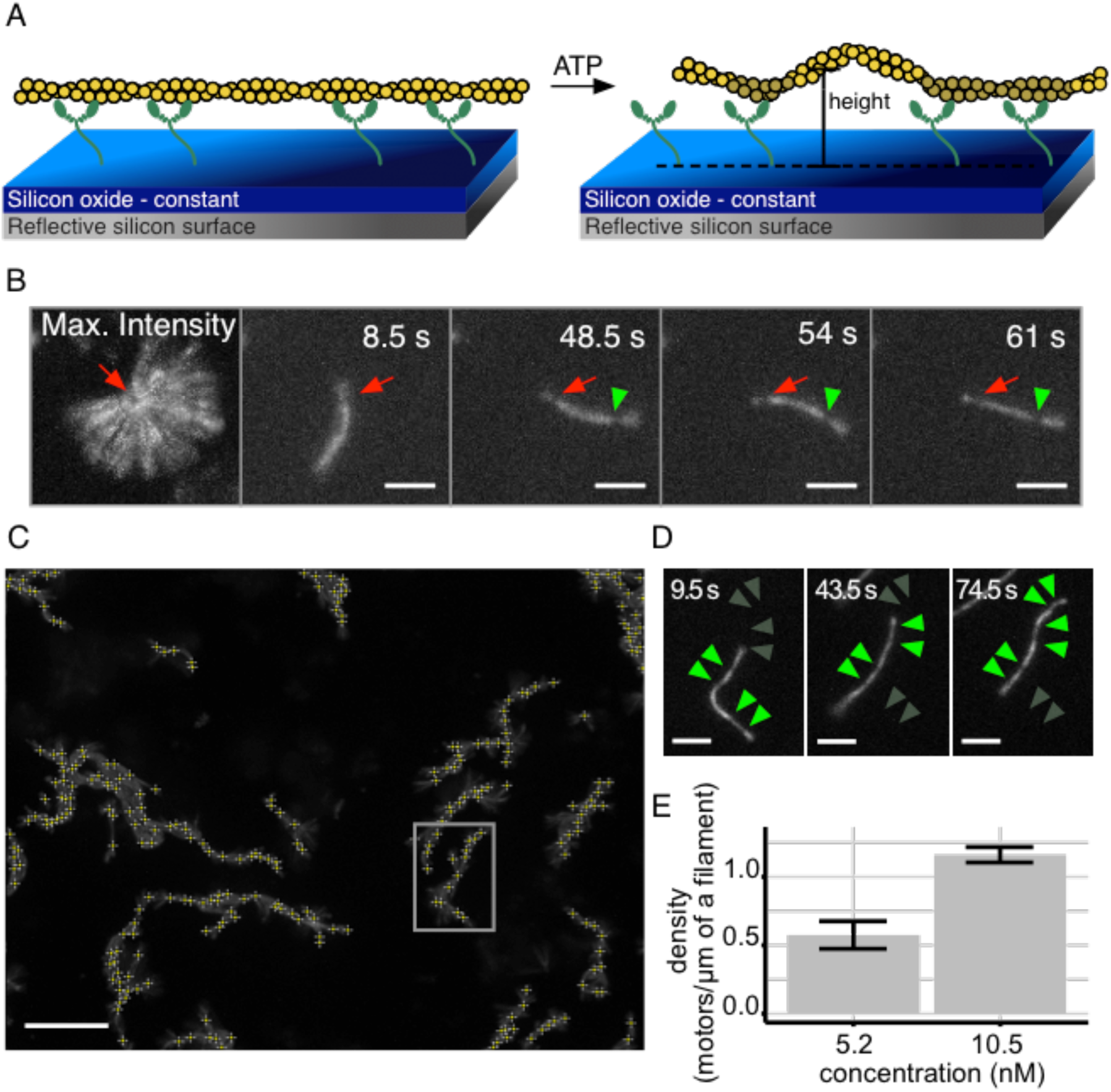
Identification of individual myosin motors in the gliding filament assay. (A) The addition of ATP allows for actin gliding. The height/intensity changes reported from now on are the consequences of actin deformation and not changes in silicon oxide thickness. (B) The dim point shows the location of myosin. The panel shows a maximum intensity projection and frames from a movie showing a single actin filament attached to the surface by a single damaged myosin (red arrow) and, in some frames (48.5 s, 54 s, 61 s), by an active motor (green arrowhead). Scale bar, 3 *μ*m. (C) FLIC robustly detects the locations of myosins in a gliding filament assay. The image shows maximum intensity projection of a single field of view of a FLIC assay. The myosins can be identified in the regions where actin was gliding. The crosses show the assigned myosin locations. The rectangle identifies the region enlarged in C. Scale bar, 10 *μ*m. (D) Example of a filament revealing the positions of 8 myosin-6s. Scale bar, 3 *μ*m. (E) The number of detected myosins is proportional to myosin concentration. The bar graph shows the surface densities of myosins derived from myosin counting and expressed as a number of myosins per 1 *μ*m of a filament. Error bars show standard error of the mean.

## THE ASSIGNMENT OF MYOSIN POSITION IN THE FLIC ASSAY

In the FLIC gliding filament assay, the addition of ATP allows the myosin to perform mechanical work. This work can result in nanoscale actin deformations such as bending the actin filament between two myosin attachment points (Figure 2A). These small changes in local actin height are detectable in the FLIC assay. The flexibility of actin filament provides a unique ability to identify the locations of myosin in the assay plane. In FLIC assay, surface-tethered myosins bind actin and bring it close to the reflective surface (Figure 2*B-D*). We can detect the location of a myosin from a short, dim segment of actin directly above the myosin. A striking example of myosin position assignment is shown in Figure 2B and Movie S1. In this sequence of images, a filament is pivoting about an attachment point: a damaged myosin-6 (red arrow). In frames 48.5 s – 61 s the filament is caught by a second, active myosin (green arrow). The active motor pulls actin into tension (straightened actin at the frame 61 s). Note that, for the rest of this manuscript, the generally rare damaged motors are excluded from the analysis.

A FLIC assay provides a robust and reliable detection method to identify and count myosins tethered to the surface, at low to intermediate myosin immobilization concentration (Figure 2C-E, Movie S2). Myosins were detected in all the regions where actin was gliding in the assay (Figure 2C). When the myosin was immobilized to the surface at half the initial protein concentration (5.2 nM versus 10.5 nM), we detected two times fewer myosins in our assay (Figure 2E). However, the relationship between myosin concentration and number of detected myosins is more complicated at higher myosin concentrations. Because of the resolution limit, the probability of undercounting the myosins increases.

## MYOSIN-POWERED ACTIN DEFORMATION

Apart from the myosin location assignment, we can identify the segments of actin under tension and compression (Figure 3). Buckled segments of actin extend higher above the surface and they appear brighter in the FLICassay (Figure 3A, segments between the yellow arrows). Occasionally, large buckles, extending beyond the first maximum of the calibration curve and out of focus can be observed, as in Figure 3A frame 71.5 s. In these large buckles, a characteristic pattern of bright edges and dim centers is visible, and the buckle appears flexible in the movie. In contrast, the segments of actin pulled into tension are brought closer to the surface and they appear dim and straight (Figure. 3A segments between the red arrows, Movie S3). We argue that this dynamic pattern of actin deformation is an outcome of asynchronous myosin stepping leading to actin tension and compression. If the filament in Figure 3B travels toward the right (blue arrow) and myosin B stochastically takes more steps than both myosin A and myosin C, then the segment of actin ahead of myosin B will be under compression and will eventually buckle. The segment of actin behind myosin B will be under tension. Therefore the pattern of buckled and taut segments of actin informs us about the relative, momentary speeds of motors.

**Figure 3.**
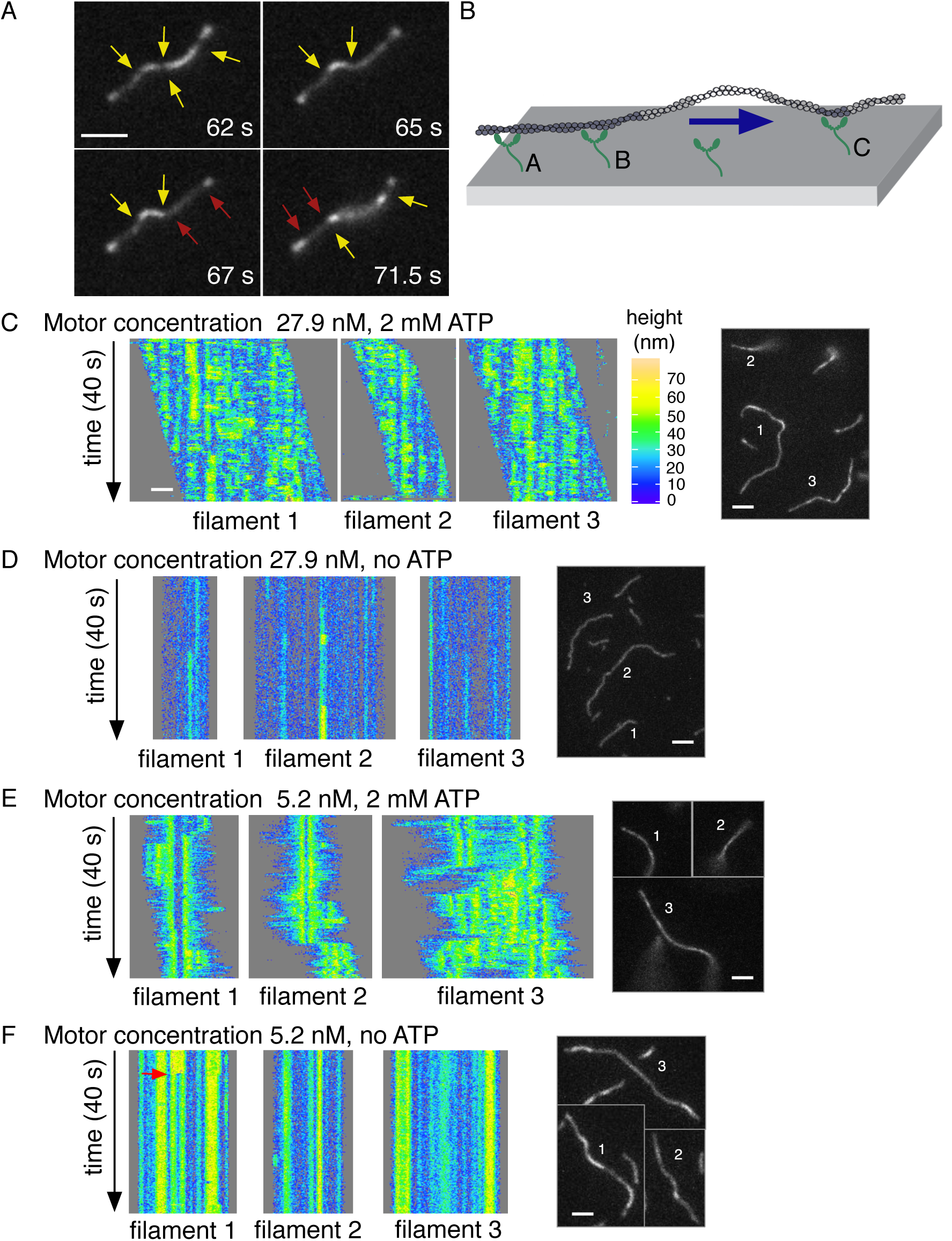
Asynchronous myosin stepping causes actin deformation. (A) FLIC reveals the state of actin between two bound motors. Frames showing the progression of a single actin filament pulled by myosin-6. Yellow arrows mark ends of actin segments buckled away from the surface between two engaged myosin-6 s. Red arrows mark ends of a segment of actin pulled into tension between two motors, and therefore pulled closer to the surface. The buckle in the last frame is large and is no longer in focus. Scale bar, 3 *μ*m. (B) The model of dynamic actin deformation. The blue arrow shows the direction of actin motion. When myosin B takes more steps than myosin C, excess actin accumulates between the two myosins, eventually forming a buckle. When myosin B takes more steps than myosins A, the intervening segment is under tension. (C-F) The magnitude of myosin-induced actin deformation. Examples of actin filaments interacting with high (C and D) and low (E and F) densities of myosin-6, in the presence (C and E) and absence (D and F) of ATP. Colors in (C) indicate the height above the surface. Notice that in (C) the filaments are moving, indicated by the diagonal band in the kymographs. Myosin attachments are visible as vertical lines corresponding to the 0-10 nm heights. The pattern of buckled segments of actin and segments of actin pulled to the surface constantly changes with ATP, and is relatively static without ATP. At low myosin concentration in (E), single myosin attachment sites are clearly visible but the ends of actin project away from the surface and out of focus. Without ATP in (F), the filaments show many static buckles, which we interpret as a consequence of sequential binding to the myosins at low surface density. Notice the myosin binding event in filament 1 (red arrow). Scale bar 3 *μ*m.

To measure the size of the buckles and to confirm their myosin-activity dependence, we scaled the fluorescence intensity of the filaments using the calibration curve shown in Figure 1B, in the presence and absence of ATP (Figure 3C-F). The myosin attachments are visible in the kymographs as vertical lines corresponding to the heights oscillating around 0 - 10 nm in Figure 3C and Figure 3E. This is consistent with the value obtained for fit parameter h = 10.4 nm, that describes the “myosin length” (see SI), in the FLIC calibration. In the presence of ATP and at the 27.9 nM myosin immobilization concentration, the buckles reach as high as ~60 nm above the surface (Figure 3C). The pattern of buckled segments of actin and segments of actin pulled toward the surface is dynamic in Figure 3C. For comparison, when the filaments from the same slide were imaged without ATP, they were pulled close to the surface over almost the entire length of the filament, without dynamic changes in filament height (Figure 3D). These results show that myosin-6 stepping causes actin buckling. Our *in vitro* assay detects these actin deformations that are mostly undetectable in standard gliding filament assays.

Interestingly, the filaments propelled by myosin at 5.2 nM immobilization concentration display a very different pattern of deformations (Figure 3E). The myosin attachment sites are very clear, however, they are so sparse that the edges of actin project away from the surface and are out of focus. The buckles are rare and they are long enough to exert a negligible force on the constraining myosins (see below). On the other hand, when only few motors are present at the surface in the absence of ATP, the filaments can be captured from the solution with buckles (Figure 3F). These buckles are static, with the exception of rare “new” binding events (Figure 3F, red arrow in filament 1).

Notice that the buckles have not been previously identified in a study of skeletal muscle myosin extension above the surface (20). We believe that the difference in myosin surface density (see later in a text) and /or the use of nonprocessive myosin-2 vs. processive myosin-6 could account for the difference between these two reports.

## BUCKLE LENGTH AND DURATION DEPEND UPON MYOSIN-6 DENSITY

To test whether the separation along the actin filament affects the collective behaviors of myosin-6, we performed the FLIC assay at different myosin immobilization concentrations. The properties of buckles vary with myosin concentration (Figure 4). At increasing myosin concentrations, the buckles are shorter (as expected when increasing the surface density of myosins) and persist for shorter amount of time (Figure. 4A). This dataset can be divided into three empirical regimes (Figure 4A, color-coded). In the first regime, “single myosins,” less than one myosin per micron of a filament is expected (see Figure 2E). Under these conditions the buckles are rarely observed, because often only a single myosin is attached. In the “dynamic buckles” regime, the surface motor density varies from 1.2 – 3.1 myosins/μm actin, using values extrapolated from the density vs. concentration relation shown in Figure 2E. Many buckles of variable size are observed at these conditions. Finally, in the “micro-buckles” regime at the highest myosin density, the buckles become smaller and persist only briefly.

**Figure 4.**
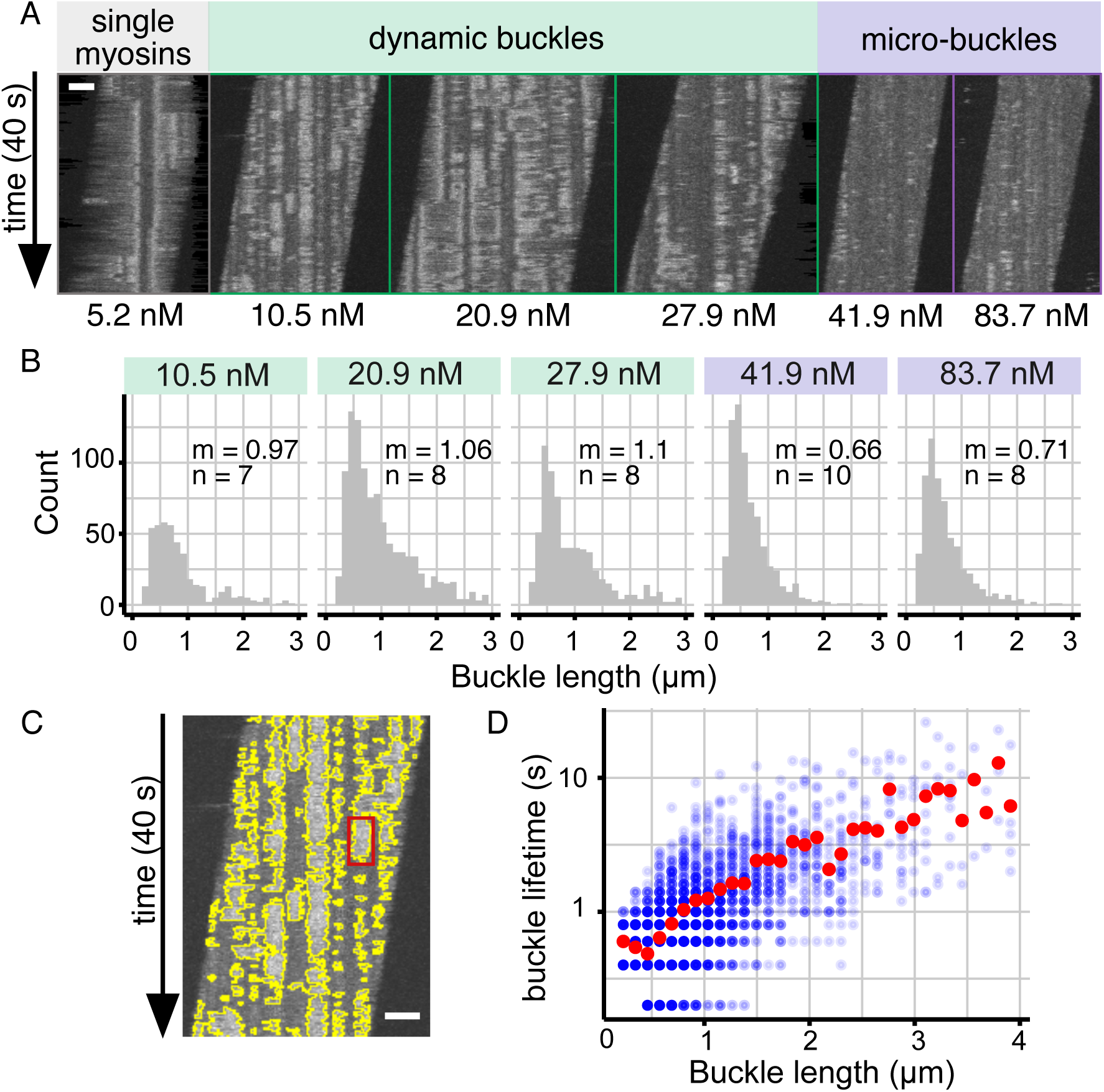
Actin filament buckling behavior depends on myosin surface density. (A) Buckles occur most robustly at intermediate myosin concentrations. The kymographs show an example filament at each myosin immobilization concentration. Three empirical regimes can be distinguished: single myosins, dynamic buckles, and micro-buckles. Scale bar, 3 *μ*m. (B) The distributions of buckle lengths show lower prevalence of very short buckles regardless of the myosin concentration. The histograms show the shift towards shorter buckles at increasing myosin concentration, however the number of detected 0.230 *μ*m long buckles is low across the entire dataset. Mean filament length: m, number of traced filaments: n. (C) Buckle lifetime and length measurements. The kymograph shows boundaries between buckled and non-buckled areas in yellow. Each buckle-containing area is then enclosed by a bounding box to determine the buckle lifetime and length (red box, see SI). Scale bar, 3 *μ*m. (D) The short buckles have a short buckle lifetime. Individual buckle lifetimes are plotted as a function of the buckle length (excluding the 5.2 nM dataset which had few buckles). The red dots show the mean buckle lifetime for every buckle length.

The distributions of buckle sizes for each immobilization concentration (excluding 5.2 nM where we do not observe frequent buckles) is shown in Figure 4B. To examine the dependence of buckle size on myosin density, the length and lifetime of the buckles was measured from the kymographs using a thresholding procedure (Figure 4C, see SI). The distributions of buckle lengths shows that increasing myosin-6 immobilization concentration reduces the number of buckles over 1 μm. However there is no dramatic increase in the number of detected small buckles (< 0.5 μm) (Figure 4B), leading to overall decrease in a buckled area of a filament. This effect is consistent with the presence of a lower limit on a buckle length, below which the filament cannot be buckled or buckles are unstable and vanish. To test this hypothesis, we examined the dependence of the buckle lifetime on the buckle length (Figure 4D). The data presented here are pooled from all myosin concentrations except for 5.2 nM myosin-6. The trend in Figure 4D shows that buckle lifetime is shorter for the short buckles.

Buckles form when the trailing myosin (myosin B in Figure 3B) moves stochastically faster than the leading myosin (myosin C in Figure 3B). Buckles disappear when the leading myosin moves faster than trailing myosin to take up the slack in the filament, or through multiple other mechanisms involving myosin attachment and detachment. As we discuss below, the force transmitted through the filament may be high enough to couple the stepping behavior of the leading and trailing myosins, or it might be inconsequential, depending on the myosin spacing. Note that these situations are quite unlike the other alternative, where the actin segment between two myosins is under tension. Under tension, the leading myosin will quickly experience stall forces because it has no available actin to translate until the trailing myosin takes a step. Thus, if the tension is maintained, the two myosins must be either coordinated or stalled. This tension-based coordination is independent of the spacing between the myosins.

The critical force for the onset of buckling of an isotropic rod can be calculated using Euler equation, and is higher for short rods (see Figure S1 and SI). When the buckled segment of actin is short, the force can be sufficiently high to retard or stall the motion of the trailing myosin. A slower trailing myosin would allow the leading myosin to catch up and retract the buckle. This mode of coordination between the motors would lead to fast retraction of short buckles, but would not affect the long buckles. The force generated by long segments of actin is too low to impact the motor kinetics. Therefore, when the long segment of actin is buckled, it is equally likely that the buckle will be extended (when the trailing myosin B steps, Figure 3B), as it is that it will be retracted (when the leading myosin C steps, Figure 3B). Thus, long segments of buckled actin filaments decouple the myosins, while short buckled segments couple the myosins.

Notice that when we describe the plausible behavior of two neighboring motors we assume an asymmetrical response to the force generated by a buckle. This assumption is consistent with what is known for myosin-5, and perhaps all other processive myosins. For myosin-5, assisting force does not increase the forward stepping rate (21, 22), while hindering force inhibits the myosin forward stepping rate (21–23).

The alternative explanation of the trend in Figure 4D would claim that shorter buckles are formed by just one or two extra steps of the trailing motor (myosin B in Figure 3B), while the longer buckles are formed by more “missteps”. If this was the case, then load-independent stepping could rapidly retract the short buckles by chance alone, because only a few (possibly just 2) subsequent steps of the leading motor are required.

## STOCHASTIC SIMULATION OF THE BUCKLE GROWTH AND RETRACTION PROCESS

To further understand the force-dependent coordination between the neighboring motors that enhances the fast retraction of short buckles, we performed stochastic simulations of a two-myosin ensemble using the Gillespie direct method (24). We ran the simulations for the three distances between the myosins, equivalent to buckle lengths, and compared the simulated results to the experimental data. The simulations were performed according to the model assuming force-dependent coordination (model 1)and unbiased stepping of myosins (model 2).

In model 1, the myosins take stochastic steps at rates determined by the forces produced by the middle segment of actin (Figure 5A). Forces are limited to tension, which switches off the leading myosin without affecting the trailing myosin, and compression, which affects the trailing myosin but not the leading. The upper limit of the compressive force experienced by the myosin is described by Euler’s critical load for buckling(Figure S1).

**Figure 5.**
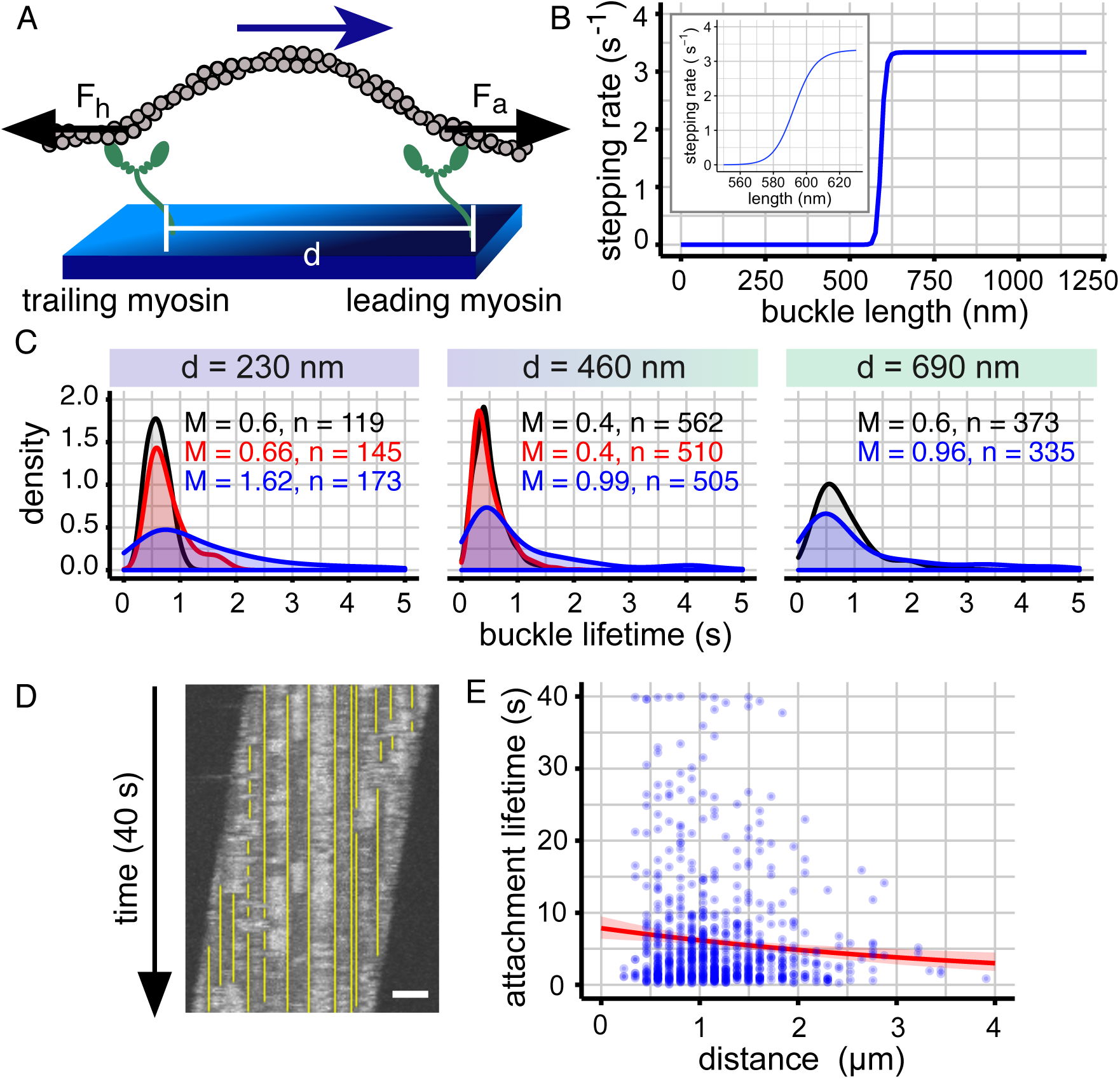
Force-based coordination mechanism limits the lifetime of short buckles and prolongs the myosin attachment lifetime. (A) Illustration of the simulation geometry. Two myosins are separated by a distance d, and are transporting an actin filament from left to right. The length of actin segment between the myosins is monitored throughout the simulation and used to determine periods of tension or compression. Hindering force, F_h_; Assisting force, F_a_. (B) The effect of buckling on the stepping rate of the left myosin, as a function of the buckle length. Stepping rates are given by Altman **(14)**, at the critical load for a buckle of the indicated length. The inset shows the vicinity of the steep transition. (C) The simulation recreates the distribution of buckle lifetimes observed in the FLIC assay for small buckles. The load-dependent model (model 1, red) reproduces the experimental lifetime distributions (black) at 230 nm and 460 nm buckles. A simple model (model 2) that allows the left myosin to step at 3.3 s^-1^ independent ofcompressive forces (blue) has a broad distribution of buckle lifetimes. Median, M; number of observations, n. (D) Myosin attachment lifetimes. An example kymograph with manually selected myosin attachment lifetimes indicated (yellow lines). Scale bar, 3 *μ*m. (E) Motors that have the close neighbors remain attached longer. The attachment lifetime data are fit to a Bell modelexponential curve 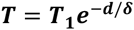, where T_1_= 7.86 and δ =4.15, by maximum likelihood estimation. The pink ribbon represents 95% confidence intervals obtained by bootstrapping 1000 datasets.

In the model 1, two myosins are separated by a constant distance *d*, and they are allowed to step forward (Figure 5A). The stepping rate of the trailing myosin, which experiences the hindering force, is modulated according to the relationship described by Altman et al. (14) (from now on called Altman regulation) and depicted in Figure 5B. The leading myosin steps at its unloaded rate ( *k* = 3.3 s^-1^), unless the actin is pulled into tension when the length of actin segment between the two myosins is shorter than the distance between them. If the tension occurs, the leading myosin stalls, as the hindering force experienced by the motor would be above the stall force (25).

To compare the simulated buckle lifetimes with my data, we set a threshold of 40 nm as a minimum height reached by the actin filament to be recognized as a buckle. To relate actin length to its height above the surface, we approximated the buckle length as an ellipse (see SI). Although this profile is not accurate, especially at the surface tangent points, the ellipse contour length falls between those of a square profile and a triangular profile and is a reasonable approximation. At all three distances between the myosins investigated in Figure 5C, a single extra step of a back motor is enough to lift actin above the 40 nm threshold (see SI) and produce a buckle that would be detected in a FLIC assay. Surprisingly, the majority of buckles observed in this dataset can be produced by a single myosin step.

Model 2 (unbiased stepping) is the same as model 1, except it does not include the Altman regulation of the forward stepping rate of the trailing myosin. Both myosins step at their unloaded rate, unless actin tension limits the leading myosin.

The 230 nm buckles are the shortest buckles we could detect in the FLIC assay. To ensure that we are not picking noise, we enforced the threshold condition in our buckle-picking procedure. The buckle has to count at least 4 pixels to be recognized. For the 230 nm buckles it means that they have to persist for at least 0.4 s (2 frames) to be recognized. We applied the same lifetime filter to the simulation of 230 nm buckles. The 460 nm buckles are the most abundant buckles in our dataset. The distance between the myosins for 460 nm buckles is small enough to experience Altman regulation of the stepping rate. The 690 nm buckles represent the situation where the force exerted by the buckle is too low to modify myosin stepping rate, therefore model 1 and 2 give the same prediction.

For myosin separation of 230 nm and 460 nm, the distribution of attachment lifetimes is well approximated by model 1 and not model 2 (Figure 5C). The model 2 creates a wide distribution of attachment lifetimes, not observed in our data for short buckles. The good agreement between experimental data and simulations according to the model 1 for 230 nm and 460 nm buckles is a strong argument in favor of force-dependent step coordination between the motors at high myosin density. When the distance between the myosins increases, the force exerted by the buckle no longer plays a role in modulating the myosin stepping rate. We would expect a good agreement between the simulations according to model 2 and the experimental data at 690 nm buckle length. However, model 2 provided a moderately good approximation for the buckle lifetime at 690 nm myosin separation. There are two main reasons to which we can attribute the difference. We might be overestimating the buckle size, because of the buckle-picking procedure (each selection is enclosed by a rectangle) and the limitations of the microscopy (diffraction limited microscopy). Another reason would be the simplicity of the simulations, as neither of our models takes under the account events like myosin backstepping, detachments, attachments of new myosins, or interactions between more than two myosins.

## ENHANCED MYOSIN-6 RUN LENGTHS AT HIGH DENSITY

The hindering force leading to myosin coordination at high density may also have an effect on myosin attachment lifetime. To investigate this possibility we traced the attachment times of the individual myosins from the kymographs (Figure 5D), which allows us to express the lifetimes as a function of the distance to the nearest neighboring myosin (Figure 5E). The results show that myosins that have close neighbors stay attached for longer. This result is consistent with the previous work showing increased dwell time of myosin-6 at high load (14). Alternatively, it can be interpreted as an avidity effect of multiple motors holding actin at a distance optimal for sustained binding.

## DISCUSSION

This work shows the application of FLIC microscopy for a study of collective myosin dynamics. In the FLIC assay, the location and attachment time of individual motors can be detected. We were able observe myosin-driven actin buckling or pulling to tension, and measure the magnitude of these actin deformations. The actin buckling is an indication of asynchronous myosin stepping. Our data show that as the myosin concentration increases the buckles become smaller and less stable. The attachment lifetime of an individual myosin also increases with nearby neighboring myosins. These two results suggest a feedback mechanism leading to myosin coordination in a gliding filament assay. Closely spaced myosins coordinate their steps to minimize the internal strain between them. Coordination is the strongest when myosins are sufficiently close, which increases the critical force required to buckle the actin filament. These results show that the mechanical coupling through actin filament regulates the collective myosin-6 behavior.

In many reports investigating the multi-motor dynamics of unconventional myosins, the myosins are linked by a quantum dot (26), DNA scaffold (7), or a vesicle (27), and they walk along surface-immobilized filament in a Total Internal Reflection (TIRF) assay. The TIRF assay enables the observation of states of the individual, labeled motors (7, 26) and mimics many aspects of cargo transport. However, the TIRF assay allows for only limited observation of actin deformation and, depending on exact experimental conditions, any actin shape variation might be severely constrained. Therefore we believe that the FLIC assay could serve as a complementary approach. It allows us to observe the dynamics of the actin filament under the influence of a fixed complement of myosin motors, which can be localized and identified as bound or detached from the actin filament.

*In vitro*, negative cooperativity, as observed by a reduced velocity and/or attenuated run length of a motor complex (relative to the model assuming no interaction between the motors), has been proposed for other processive motors including myosin 5 (5, 7, 28). In our assay, we observe an increased attachment lifetime of an individual myosin if another myosin is in its vicinity (Figure 5E). This response is consistent with increased dwell time of myosin-6 when external loads are applied (14), however, it is unclear to what extent this effect depends upon the experimental geometry.

In a FLIC assay, myosins are mechanically coupled through the actin filament. It is expected that the elastic properties of F-actin (e.g., the persistence length, Lp, used to calculate the Euler force) play crucial role in modulating the behavior of the myosin ensemble. The role of filament elasticity is in agreement with previous reports, which found that the behavior of a motor assembly depends upon the properties of the motors and the connections between them (5, 7, 27, 29).

The data presented here show robust actin buckling at the intermediate concentrations of myosin-6. Actin buckling powered by myosin-2 is crucial for symmetry breaking between tensile and contractile forces, which is necessary to drive network contraction in a minimal model of the cell actomyosin cortex (30, 31). We speculate that myosin-6 motors attached to a large cellular structures that provide the necessary separation, like the Golgi apparatus or the plasma membrane between the stereocilia (12, 13, 32), could buckle a segment of actin between them. However, the force transmitted between the myosins in this scenario is expected to be low and insufficient to lead to the coordination of myosin stepping. Additionally, membrane fluidity would play role in modulating the extent of myosin-6-driven actin deformation. On the other hand, when myosin-6 transports smaller objects like clathrin-coated vesicles that are only 100 – 150 nm in diameter (1), the myosins would be mechanically coupled through the filament. If the buckling occurred between the myosin-6s attached to endocytic vesicles, it would still transmit a high force between the myosins and would lead to myosin coordination. The buckling could relax the normal force experienced by the myosin-6s attached to the curved vesicle.

The study of myosin-6 collective behavior presented here shows the unique utility of the FLIC assay to investigate the actomyosin interaction. However there are still many questions that remain to be addressed. Apart from coordination of a forward step between neighboring motors, does the decreased buckling at high myosin concentration (see Figure 4A) indicate “lock-step” type of synchronization between myosins? Such synchronization between motors has recently been proposed as a possible mode of motor cooperation (29). Moreover, how do different myosin classes behave in a FLIC assay? These and many other questions should be the subject of future studies.

## MATERIALS AND METHODS

### MYOSIN CONSTRUCTS AND PROTEIN REAGENTS

The myosin-6-HMM-GCN4-Flag-Ctag construct and myosin-6-HMM-GCN4-Flag-YFP construct (used only for data presented in Figure 2B and 3A) contained porcine myosin 6 (GenBank accession number XP_005659483, amino acids 1-994). Myosin-6-Ctag construct was expressed using recombinant baculovirus in Sf9 insect cells. Myosin-6-YFP construct was expressed in Sf9 insect cells using the Insect Direct expression system. All the myosin-6 constructs were co-expressed with baculovirus containing calmodulin and the light-chain MLC-1sa from human.

Affinity clamp protein was expressed and purified from bacterial culture as previously described (33, 34). Calmodulin was expressed and purified from bacterial culture as previously described (35). Actin was purified from chicken breasts as previously described in and polymerized to F-actin at 10 μM monomer concentration as described in (36). Here, 1% of biotinylated actin monomers (labeled at Cys 374 by biotin-maleimide (37,38)) was used.

### MYOSIN PURIFICATION

Myosin-6 was purified using Flag affinity chromatography. Briefly, Sf9 cells were infected with the appropriate baculovirus and cultured for 48h. Cells were harvested by centrifugation and resuspended in lysis buffer (50 mM Tris pH 7.7, 150 mM KCl, 4 mM MgCl_2_, 0.5 mM EDTA, 1 mM EGTA, 0.1% Triton X-100, 7% sucrose, 2 mM ATP and protease inhibitors). The cells were freeze-thawed and nutated for 30 minutes at 4°C to allow the myosin to dissociate from actin. The lysate was spun down and the supernatant was incubated with the anti-Flag resin (Sigma) for 1 hour at 4°C with nutation. The resin was pulled down by either light spin or gravitationally and resuspended in wash buffer (20 mM imidazole pH 7.5, 150 mM KCl, 5 mM MgCl2, 1 mM EDTA, 1 mM EGTA, 3 mM ATP, 0.5 mM DTT and protease inhibitors). The Pierce drip column was assembled and the anti-Flag resin was transferred to the column. The resin was extensively washed and then the column was capped and incubated for 1 hour at 4°C with a wash buffer containing 0.2 mg/ml Flag peptide. The elution was collected and dialyzed over night against dialysis buffer (25 mM imidazole pH 7.5, 1 mM EGTA, 4 mM MgCl2, 150 mM KCl, 50% glycerol, 1 mM DTT). The myosins were stored at -20 ^°^C.

### SILICON WAFER PREPARATION

Mechanical grade silicon wafers coated with a 1-1.7 nm thick layer of silicon oxide were cleaned extensively with acetone and methanol. The additional layer of silicon oxide was deposited, to the desired final thickness reported in the main text, using AJA Orion 5 UHV sputtering system. The thickness of silicon oxide layer was measured by ellipsometry using a Gaertner Waferskan Ellipsometer. Between assays, the silicon wafers were cleaned by piranha solution.

### FLIC ASSAY

The flow cell was constructed from silicon wafer coated with ~20 nm SiO2 layer (except for data presented in Figure 2B and 3A, where the SiO^2^ thickness was ~1.7 nm), two pieces of double-sided tape, and glass coverslip. Flow cells were incubated with affinity clamp protein (0.33 mg/ml in phosphate-buffered saline pH 7.3: 137 mM NaCl, 2.7 mM KCl, 4.3 mM Na_2_HPO_4_•7H2O, 1.4 mM KH_2_PO_4_; 2 min) or with anti-GFP (0.05 mg/ml in PBS), followed by a bovine serum albumin block (1 mg/ml in Assay Buffer, AB, 2 min). The composition of AB buffer was: 25 mM imidazole, pH 7.5, 25 mM KCl, 1 mM EGTA, 4 mM MgCl2, 10 mM dithiothreitol. Myosin 6 was added to the flow cell at the concentrations indicated in the main text (10 μl of the myosin diluted in AB, 2 min). The flow cell was rinsed extensively with AB. Then, 20 nM F-actin stabilized with tetramethylrhodamine-phalloidin (Sigma) or Atto647N-phalloidin (ATTO-TEC, data presented in Figure 2B, 3A) was added and incubated for 2 min. The flow cell was washed and static filaments were imaged in the AB buffer containing 0.086 mg/ml glucose oxidase, 0.014 mg/ml catalase, and 0.09 mg/ml glucose. For moving filaments, the motility buffer (2 mM ATP, 0.086 mg/ml glucose oxidase, 0.014 mg/ml catalase, and 0.09 mg/ml glucose in AB) was added. Imaging was performed using a Zeiss Axiovert 200 microscope with an Andor Luca camera and Olympus 63x 1.2 water immersion objective.

See the Supporting Information for further experimental and analysis procedures.

## AUTHOR CONTRIBUTIONS

AKK and RSR designed research; AKK performed research; JJR contributed reagents; AKK and RSR analyzed data and wrote the manuscript.

## ACKNOWLEDGMENTS

The authors thank Sally Horne-Badovinac, Tobin Sosnick, Margaret Gardel, Kim Weirich and Patrick McCall for helpful discussions, and members of the Rock lab for support. This work was supported by NIH R01s GM078450 and GM109863 (to R.S.R.). We acknowledge the University of Chicago Searle Cleanroom and Nanofabrication Facility for support of this work.

